# A new function for Prokineticin 2: recruitment of SVZ-derived neuroblasts to the injured cortex in a mouse model of traumatic brain injury

**DOI:** 10.1101/273698

**Authors:** Mayara Vieira Mundim, Laura Nicoleti Zamproni, Agnes Araújo Sardinha Pinto, Layla Testa Galindo, André Machado Xavier, Isaias Glezer, Marimélia Porcionatto

## Abstract

Traumatic brain injury is an important cause of mortality and morbidity all over the world. After the initial injury there is a cascade of cellular and molecular events that ultimately lead to cell death. Therapies aim not only to counteract these mechanisms but also to replenish the lost cell population in order to achieve a better recovery. The adult mammal brain in not as plastic as the postnatal, but it has at least two neurogenic regions that maintains physiological functions in the brain; the subgranular zone of the dentate gyrus of the hippocampus, which produces neurons that integrate locally, and the subventricular zone (SVZ) of the lateral ventricles, that produces neuroblasts that migrate through the rostral migratory stream (RMS) to the olfactory bulbs. Brain injuries, as well as neurodegenerative diseases, induce the SVZ to respond by increasing cell proliferation and migration to the injured areas. Here we report that SVZ cells migrate to the injured cortex after traumatic brain injury in mice, and that the physiological RMS migration is not impaired. We also show that Prokineticin 2 (PROK2), a chemokine important for the olfactory bulb neurogenesis by promoting the directional migration of neuroblasts, is induced in the injured cortex. Using PROK2 receptor antagonist and recombinant PROK2 we show for the first time that PROK2 can directionally attract SVZ cells *in vitro* and *in vivo*. The data we present here links one more element of the inflammatory process, PROK2 secreted by microglia, to the attempt to regenerate an acutely injured mammalian cortex.

**Abbreviations:**

SGZ: subgranular zone
SVZ: subventricular zone
RMS: rostral migratory stream
PROK2: Prokineticin 2

## Introduction

Traumatic brain injury is an important cause of morbidity and mortality all over the world. In the United States, CDC estimated that in 2010 approximately 2.5 million people were assisted in hospitals after a TBI event. Of those, approximately 2% died. The incidence data of TBI-related disabilities is limited, but extrapolation of two US states data estimates that 3.2 million-5.3 million people live with TBI-related disabilities in the US. Those include cognitive, behavior, motor and somatic symptoms, that not only costs millions to the state but also represents a decrease in the quality of life of those injured and their families (CDC, 2015).

The pathophysiology of TBI includes primary and secondary injuries; the primary injury is the direct result of the physical trauma, and the latter develops into a cascade of cellular and molecular degenerative mechanisms that accounts for the main outcome of brain injury and is the target of therapies (Galgano et al., 2016).

Since the discovery that the adult mammal brain is not quiescent (Altman, 1963), and has neurogenic niches that can respond to brain pathologies by cell proliferation and migration to injured areas, research has been conducted with the main focus on cell-based therapies (Altman, 1963; Chang et al., 2016; Dizon et al., 2010; Eriksson et al., 1998; Harting et al., 2009; Park et al., 2012; Reis et al., 2017; Reynolds & Weiss, 1992; Sun et al., 2011).

The main neurogenic areas in the adult brain are the subgranular zone (SGZ) of the hippocampus dentate gyrus, and the subventricular zone (SVZ) outlining the lateral ventricles. The neural stem cells in the SVZ give rise to neuroblasts that physiologically migrate through the rostral migratory stream RMS) to the olfactory bulbs where they differentiate into olfactory neurons (Alvarez-Buylla & Garcia-Verdugo, 2002; Gage, 2000; Lois & Alvarez-Buylla, 1994). Prokineticin 2 (PROK2) is a chemokine involved in the olfactory bulb neurogenesis, and is responsible for directing neuroblasts migration (Ng et al., 2005). PROK2 activates two G-protein coupled receptors, PKR1 and PKR2, that are differentially expressed throughout the brain, conferring a broad spectrum of functions, such as regulation of circadian rhythm, thermoregulation and energy expenditure (Cheng et al., 2002; Zhou et al., 2012). Recently it has been reported that PROK2 is upregulated in some brain pathologies such as Parkinson’s and Alzheimer’s disease and stroke (Cheng et al., 2012; Gordon et al., 2016; Severini et al., 2015). Here we investigated the role of PROK2 in a traumatic brain injury model. We show, for the first time, that PROK2 is expressed by microglia at the injured site as early as one day after injury, and that it enhances migration of SVZ-born neuroblasts to the injured area.

## Material and methods

### Animals

Male C57BL/6 mice aged 2 days, 6 or 12 weeks used for experiments were maintained under a 12 hours light/dark cycle with access to water and food *ad libitum*. All experimental procedures and animal handling were performed in accordance with the UK Animals (Scientific Procedures) Act of 1986 Home Office regulations and approved by the Committee for Ethics in the Use of Animals from Universidade Federal de São Paulo (CEUA 747918).

### Surgical procedures

Mice were anesthetized with xylazine (Ceva) 10 mg/kg of body weight, ketamine (Syntec) 100 mg/kg and acepromazine (Vetnil) 1 mg/kg and placed in a stereotaxic apparatus. A blunt metal needle was inserted three times into the mice primary motor cortex [AP 0mm, ML +1 mm, DV -0.75 mm (Paxinos & Franklin, 2001)]. After the lesion the skin was stapled and the mice were returned to their cages for recovery. The animals received analgesic (acethaminophen 1mg/mL, EMS) in the drinking water for two days after the surgery.

### BrdU and EdU administration

For proliferation studies two protocols of BrdU injections were used: one injection of BrdU (Sigma, 50 mg/kg of body weight, in 0.9 % NaCl solution) 30 minutes after the injury, and then two more injections every 2 hours; and one injection 6 hours after the injury and then 4 more injections every 24 hours. For migration studies mice were injected with 4 doses of BrdU every 12 hours with a washout period of 15 hours before the injury or recombinant PROK2 administration, and EdU (Invitrogen) was injected in a single dose (50 mg/kg body weight) 4 hours after the injury. The protocols used are indicated in each experiment.

### PKRA7 and PROK2 administration

The PROK2 receptor antagonist PKRA7 (20mg/kg, Merck) was administered i.p. 1 hour after the injury and then every 24 hours for 3 days. Control animals were injected with 10 % DMSO. A single dose of recombinant PROK2 (10 pMol, Peprotech, USA) was administered in the motor cortex of mice [AP 0mm, ML +1 mm, DV -0.75 mm (Paxinos & Franklin, 2001)]. Control animals were injected with 0.9 % NaCl solution.

### Immunohistochemistry and image acquisition

On the last day of the experiment (indicated in each experiment), mice were anesthetized and intracardially perfused with ice-cold paraformaldehyde (4 % PFA) in 0.1M PBS. The brains were removed, postfixed in 4 % PFA overnight at 4 °C, and cryopreserved in 30% sucrose solution at 4 °C for 48 hours. The brains were frozen on dry ice. Using a Leica CM1850 cryostat 30 μm serial tissue sections were cut with 120 μm interval for RMS and 360 μm for the SVZ sections. Free-floating sections were washed in 0.01 M phosphate buffer saline and treated with 50 mM Glycine for 15 minutes. Blocking was done with 5 % normal goat serum and 0.1 % Triton X-100 for 1 hour. Primary antibodies were diluted in blocking solution and incubated on sections overnight at 4 °C. Secondary antibodies were diluted in blocking solution and DAPI (Sigma, 1:1000) and incubated in sections for 1 hour at room temperature. Sections were washed with phosphate buffer and mounted using fluoromount (Electron Microscopy Sciences). For BrdU staining the sections were treated with 2 N HCl for 30 minutes at 37 °C after the initial wash, and were not colored with DAPI. EdU was detected using the Click-iT™ EdU Alexa Fluor™ 488 Imaging Kit (Molecular Probes) according to the manufacture’s instruction. The following primary antibodies were used: rat anti-BrdU (1:500, Bio-Rad OBT0030), guinea pig anti-Dcx (1:500, Millipore, ab2253), chicken anti-GFAP (1:500, Millipore ab5541), rabbit anti-PROK2 (1:200, Abcam ab76747), mouse anti-CD45 (1:100, Affymetrix 48-0451-80), goat anti-Iba1 (1:400, Abcam ab5076). Alexa Fluor™ secondary antibodies (1:500, Molecular Probes) were used. A Zeiss 710 confocal microscope was used for image acquisition. Images were acquired from 3 sections per animal, and 15 μm with 1.5 μm intervals per section were imaged and used for analysis and cell counting. Fiji was used for image analysis; cells were counted using the Cell Counter Plugin (Schindelin et al., 2012).

### qPCR

A chunk of the injured cortical area and both olfactory bulbs were used for RNA extraction with Trizol^®^ (Invitrogen) and quantified using the spectrophotometer NanoVue Plus (GE Healthcare). One μg of total RNA was reverse-transcribed with Oligo(dT)15 Primer (Promega) and ImProm-II Reverse Transcription System (Promega) with Recombinant RNasin^®^ Ribonuclease inhibitor (Promega) and MgCl2. qPCR was performed using 20 ng cDNA and Brilliant® II SYBR® Green QPCR Master Mix (Stratagene), in an Applied Biosystems® 7500 Real-Time PCR System. Thermal cycling was 95 °C for 10 minutes, 40 × 95 °C for 15 seconds, 60 °C for 1 minute and 72 °C for 30 seconds. Dissociation curve was done at 95 °C for 1 minute, 60 °C for 30 seconds and 95 °C for 30 seconds. The primers used were: *Prok2* – sense 5’-CGGAGGATGCACCACACCT-3’ and antisense 5’-TTTCCGGGCCAAGCAAATAAACCG-3’, *Cxcl12* – sense 5’-ATCCTCAACACTCCAAACTGTGCC-3’, and antisense 5’-TTCAGACCTAGGCTCCTCCTGTAA-3’, *Hprt* – sense 5’-CTCATGGACTGATTATGGACAGGAC-3’ and antisense 5’-GCAGGTCAGCAAAGAACTTATAGCC-3’.

### Plasmids

Murine *Prok2* coding sequence was amplified from the cDNA of cortical chunks from five injured mice by PCR using *Pfu* DNA polymerase, Recombinant (Thermo Fisher Scientific, cat#EP0501) according to manufacturer instructions. The primers for *Prok2* were: forward 5’-**GGAATTCC**ATGGGGGACCCGCGCT-3’ and reverse 5’-**CGGGATCCC**CATTTCCGGGCCAAGCAAA-3’; the restriction sites *Eco*RI and *Bam*HI, respectively, are indicated. The PCR product was digested with the indicated restriction enzymes and cloned into pEGFP-N3 (Clontech) mammalian expression vector. For the control empty vector, *Eco*RI/*Bam*HI digested plasmid was treated with blunting enzyme from CloneJET PCR cloning kit (Thermo Fischer Scientific, cat#K1239) and re-ligated with T4 DNA ligase (NEB, cat#M0202V). The plasmids were expanded in DH5α *E. Coli* and purified with maxi-prep kit according to the supplier’s protocol (Qiagen, Valencia, CA, USA). The constructs were verified by restriction analysis and DNA sequencing with BigDye Terminator v3.1 Cycle Sequencing kit (Applied, cat#4334755).

### Transfection of HEK293T cells

Cells were cultured in monolayer in DMEM High Glucose medium (Gibco), 10 % fetal bovine serum (Cultilab), 1 % glutamine (Sigma) and 1 % Penicillin/Streptomycin (Gibco) at 37 °C and 5 % CO2. When 70 % confluent cells were transfected with pEGFP-PK2 or pEGFP plasmids using the Calcium Phosphate method. Briefly, 15 μg DNA, 2 M CaCl2 and H2O were mixed in a vial and 2x HBS was added after. The mix was gently dropped around the plate and the cells were incubated for 24 hours at 37 °C and 5 % CO2. The medium was changed after that and cells were incubated for another 24 hours. After that the cells were trypsinized and cultured as hanging drops in a density of 2.5×10^6^/mL. HEK293T spheroids were allowed to form for 1 day.

### Neural stem cell culture

Neural stem cells were obtained from 6 weeks old C57BL/6 mice. Briefly, mice were euthanized and their brains were removed. The SVZ was dissected and the cells were dissociated by incubation with Accutase (Gibco) for 5 minutes at 37 °C. After mechanical dissociation cells were strained in a 40 μm mesh (BD Biosciences) and plated in Poly-HEMA (Sigma) precoated flasks, in complete medium containing DMEM/F12 1:1 (Gibco), 2 % B27 supplement (Gibco), 20 ng/ml EGF (Sigma), 20 ng/ml FGF2 (R&D Systems), 1 % penicillin/streptomycin (Gibco), and 5 μg/mL heparin (Sigma). After neurosphere formation (100-150 μm) they were passaged and allowed to form once again, and used after third passage.

### Co-culture

Transfected HEK293T spheroids and neurospheres were plated very closely in a 20 μL drop of 6mg/mL matrigel (BD). One spheroid and one neurosphere were plated in each well. The wells were filled with complete medium without growth factors and incubated at 37 °C and 5 % CO2 for 5 days.

### Astrocyte culture

Cells were extracted from 2 days old C57BL/6 mice. After euthanasia the cortices were dissected, chopped and washed in a Ca^2+^ e Mg^2+^ free Hanks solution containing 10 % bovine serum and 1 % penicillin/streptomycin, and in a Versene solution. The tissue was incubated with 0.25 % trypsin (Gibco) and 1mM EDTA for 30 minutes at 37 °C and mechanically dissociated. After washes the cells were resuspended and cultured as monolayer in medium containing DMEM/F12 1:1, 10 % inactivated bovine serum and 1 % penicillin/streptomycin. Twenty-four hours later medium was changed and then every 48 hours. For qPCR experiments astrocytes RNA was extracted using the PureLink RNA Micro Kit (Invitrogen) according to manufacturer’s protocol. cDNA synthesis and qPCR were done as described for qPCR. For quantitative experiments, proteins were extracted using RIPA buffer and ELISA was performed using Duoset dy460 ELISA Kit (Red Systems) following manufacturer’s protocol.

### Statistics

Statistical analysis was done using GraphPad Prism. The graphs presented show the mean ± standard error. The difference between groups was assessed by Student’s t-test, unless stated otherwise. Groups were considered statistically different when p≤0.05.

## Results and Discussion

### Cortical injury induces astrogliosis and cell proliferation in the cortex and SVZ

We used a model of brain injury caused by a punctual wound, which simulates an open head injury (Fig. 1C). We show that in this model intense astrogliosis in the SVZ, corpus callosum and cortex appears as early as 12 hours after the injury (Fig. 1E). We administered BrdU 30 minutes after the injury and then every 2 hours twice to label cells proliferating in response to the injury (Fig. 1A). We observed cell proliferation in the SVZ (Fig 1D-F), but did not find many dividing cells in the cortex near the injury in the period of 18 hours post injury (data not shown). We then administered BrdU to the animals beginning 6 hours after the injury and then every 24 hours for 4 days (Fig. 1B). On the fifth day many BrdU+ cells were found in the cortex, mainly concentrated near injury site (Fig. 1G), and proliferating astrocytes were among the BrdU+ cells in the cortex (Fig. 1H). Proliferating reactive astrocytes in lesion areas were reported to have the capacity to form neurospheres with self-renewing and multipotent properties *in vitro* (Buffo et al., 2010; Lang et al., 2004). This stem cell-like response of the astrocytes was also seen *in vivo* (Sirko et al., 2015), and proposes another role for astrocytes other than brain housekeepers. Our model recapitulates major events after injury and is suitable to evaluate the cellular and molecular responses after a traumatic brain injury.

**Figure 1.**
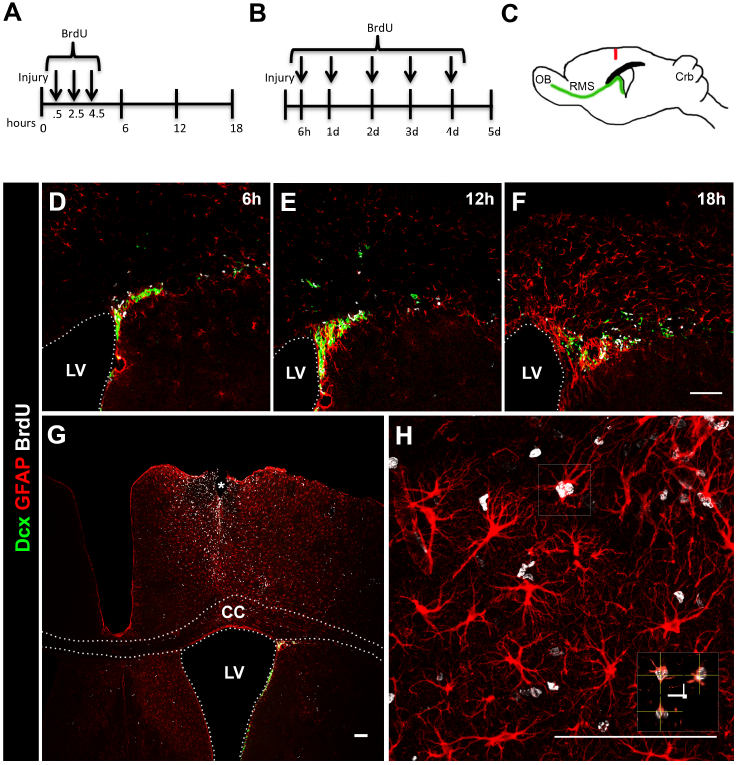
Cortical injury induces astrogliosis in the cortex and SVZ. (A,B) BrdU administration protocols. (C) Drawing of a sagittal mouse brain TBI model showing the location of needle insertion. (D-F) 6, 12 and 18 hours, and (G-H) 5 days after TBI. Neuroblasts (Dcx, green), proliferating cells (BrdU, white) and reactive astrocytes (GFAP, red) in the dorso-lateral SVZ, corpus callosum and cortex. LV, lateral ventricle; CC, corpus callosum. Scale bar: 100mm.

### SVZ-derived neuroblasts migrate to the cortex after cortical injury

In order to evaluate the migration of SVZ cells towards the injured cortex, we administered BrdU to the animals every 12 hours for two days prior to the injury (Fig. 2A). An undamaged brain contains only two regions with constant cell proliferation, the neurogenic SVZ and SGZ, therefore any BrdU+cells found in the brain are mainly derived from these regions. To label the cells that proliferated in response to the injury we administered EdU to the animals 4 hours after performing the injury (Fig. 2A). The administration of the thymidine analog BrdU up to 4.5 hours after the injury does not label cortical cells (Fig. 1), so any EdU+ cell would be derived from the neurogenic regions. The SGZ produces cells that integrate the hippocampus and there is no demonstration in the literature that these cells migrate to the injured cortex. That said, we found many SVZ derived BrdU+ and EdU+ cells leaving the SVZ, in the corpus callosum, cortex, and mainly near the site of injury (Fig. 2C), indicating that not only the cells that proliferate in response to the injury, but also the ones that are physiologically proliferating in the SVZ migrate to injured areas. Some of the cells we found were BrdU+EdU+, indicating that a portion of the SVZ cells underwent cell cycle re-entry after the injury and migrated to the cortex. BrdU+ cells did not appear to migrate only from the SVZ, but some SVZ-derived cells that were already on the rostral migratory stream on route to the olfactory bulb seemed to deviate from their original course and migrate towards the injured cortex (Fig. 2D), and many neuroblasts were found near the site of the injury (Fig. 2E, F). In the inner cortical layers, in proximity to the corpus callosum, the neuroblasts presented a migratory morphology, and differently from the RMS, they were migrating as single cells (Fig. 2E). However, the neuroblasts seemed to be migrating following a route; there are descriptions of neuroblasts migrating using blood vessels and/or reactive astrocytes as scaffold and/or support (Saha et al., 2013). Close to the site of the injury the neuroblasts presented smaller cell body and increased complexity of dendrites (Fig. 2F, red Dcx+ cells), probably in the attempt to integrate the injured area and make new synaptic connections during the early post injury period.

**Figure 2.**
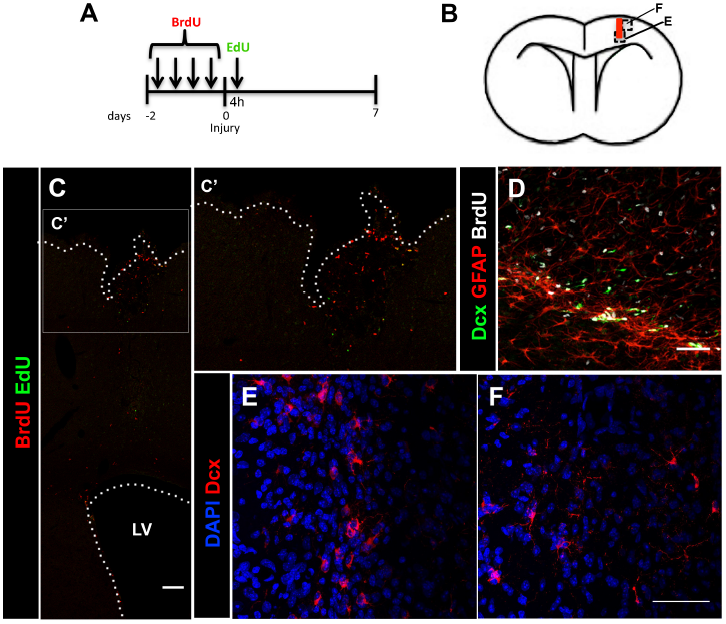
Cells migrate out of the SVZ towards the cortical injury. (A) BrdU and EdU administration protocol. (B) Drawing of a coronal mouse brain TBI model showing the location of needle insertion, and indication of areas shown in panels E and F. (C) BrdU+ (red) and EdU+ (green) cells migrated from the SVZ into the injured cortex area after 7 days. (C’) Higher magnification of the injured cortex showed in C. (D) BrdU+ cells (white) and neuroblasts (Dcx, green) emigrate from the RMS after TBI. (E,F) Neuroblasts (Dcx, red) migrate from the SVZ to the injured cortex area. LV, lateral ventricle. Scale bar: (C) 100μm and (D-F) 50μm.

### Cortical injury does not impair migration of SVZ cells to the olfactory bulb

Our protocol for BrdU and EdU incorporation allows us to differentiate between neuroblasts generated before and after the injury. BrdU+ cells are the ones originated by the continuous proliferation of SVZ neural stem cells before the injury and EdU+ cells are generated in response to the injury (Fig. 3A). We found that fewer BrdU+ cells remained in the SVZ 7 days after the injury (Fig. 3D, E), but there was no difference in the number of EdU+ cells between the groups (Fig. 3F). Also we found no difference in the number of BrdU+EdU+ cells (Fig. 3G) indicating no changes in the cell cycle re-entry index. The percentage of BrdU+ cells that were Dcx+ was not altered by the injury (Fig. 3H), but more EdU+ cells were also Dcx+ after the injury (Fig. 3I). These data indicate that more SVZ cells generated after the injury produced neuroblasts (Dcx+EdU+ cells) that emigrated from the SVZ. Brain injuries increase cell proliferation in the SVZ (Radomski et al., 2013; Saha et al., 2013; Theus et al., 2010), but since we did not see more EdU+ cells in the SVZ, and the fact that more Dcx+EdU+ cells were found in the SVZ after the injury we hypothesized that a increased number of cells that proliferated after the injury differentiated into neuroblasts probably in order to migrate to the injured area.

**Figure 3.**
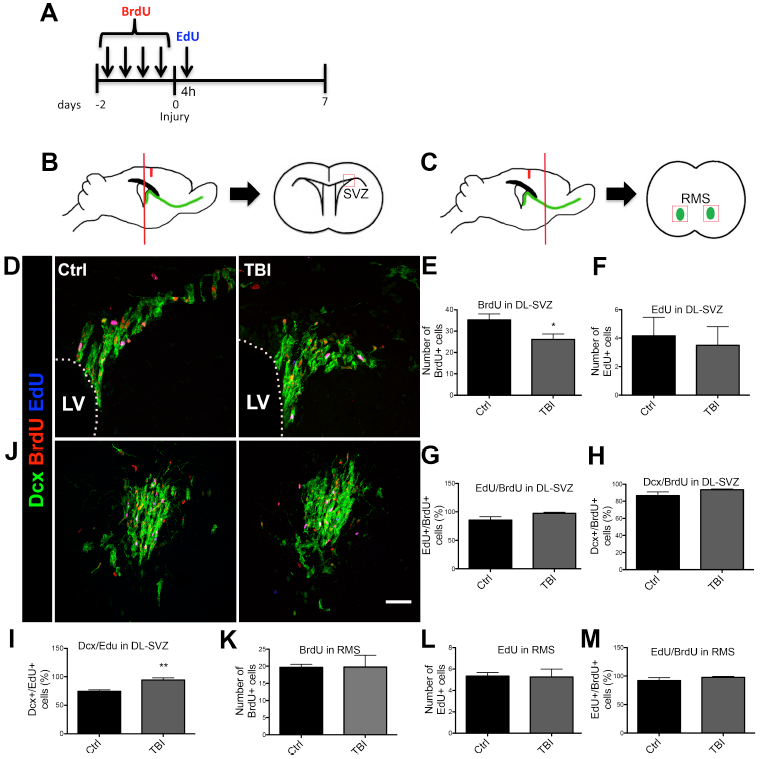
Cortical injury does not impair migration of SVZ-born neuroblasts to the olfactory bulb through the RMS. (A) BrdU and EdU administration protocol. (B-C) Drawing of sagittal and coronal mouse brain showing the areas corresponding to the sections used to capture images of the SVZ and RMS. (D-I) Fewer BrdU+ cells and more Edu+Dcx+ are found in the SVZ 7 days after TBI. (J) The number of BrdU+ and Edu+ cells in the RMS is not changed 7 days after TBI. LV, lateral ventricle; DL-SVZ, dorsolateral SVZ; RMS, rostral migratory stream. Graphs show mean ± SEM. Scale bar: 50μm.

What happens to the physiological migration of SVZ-derived neural stem cells after brain injuries is still controversial in the literature. The double labeling protocol using thymidine analogs incorporation to follow SVZ-derived cells is broadly used to trace the cells and find their final destination after injury. However, the results largely depend on the incorporation protocol that was used, and since most of the experiments are of end point type, the results also depend on how many days after the injury the brain was analyzed. Here we show that the number of BrdU+ and EdU+ cells in the RMS was not altered 7 days after the injury (Fig. 3K-M). We found some BrdU+ and EdU+ cells out of the RMS in the injured group, and we also showed that some cells migrate out of the RMS after the injury (Fig. 2D), but we hypothesize that the lack of alteration in the RMS migration is due to increased proliferation in the SVZ, thus compensating for the normal migration of neuroblasts towards the olfactory bulb.

Many soluble molecules produced and secreted at a cortical acute injury site have been reported, especially chemokines derived from immune cells and astrocytes CXCL12 is one of the chemokines which expression is increased after acute injury to the brain and that attracts neuroblasts to an injury site (Dalgard et al., 2012; Galindo et al., 2011). We were interested in knowing if the migrating neuroblasts responded differently to two distinct chemotactic stimuli, CXCL12, as found at the injury site, and PROK2, constitutively expressed in the olfactory bulb and considered one of the key chemoattractants to direct neuroblasts that arrive to the olfactory bulb (Ng et al., 2005).

### Prokineticin 2 is expressed in the injured cortex

PROK2 is mostly expressed in the granular and periglomerular layers of the olfactory bulb, exerting its function in neurogenesis, and also has a discrete expression throughout the normal brain (Cheng et al., 2006). Since to our knowledge there was no information regarding the expression of PROK2 following brain injury, we investigated *Prok2* mRNA expression in the olfactory bulb in different time points after the TBI model (Fig. 4A). We found that *Prok2* expression decreased one day after the injury, but after the second day, expression was more than twofold. After four days the expression returned to basal level and remained for at least 14 days. The sudden decrease in *Prok2* expression could be a general response to injuries, since a similar injury was performed in the cerebellum also caused decrease in *Prok2* mRNA expression (data not shown). The following rapid overexpression could be a rebound, probably to prevent further consequences to the RMS migration that could have happened by the diminished *Prok2* in the olfactory bulb.

**Figure 4.**
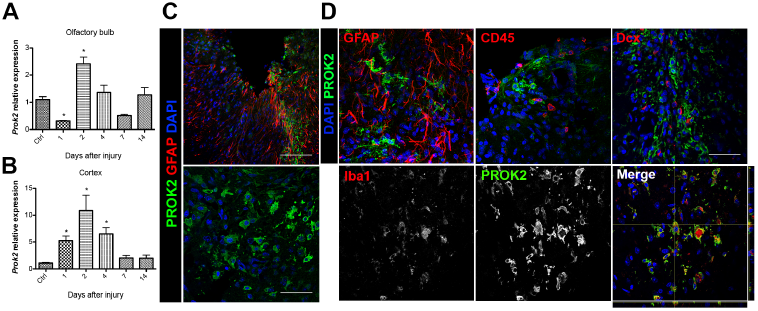
Prokineticin 2 is expressed by microglia in the injured cortex. (A, B) Time course expression of *Prok2* mRNA after TBI in the olfactory bulb and cortex. Expression was normalized to *hprt*. (C) PROK2 (green) is expressed in the injured cortex. Graphs show mean ± SEM. Scale bar 100μm (top image) and 50μm (bottom image). (D) PROK2 (green) is expressed exclusively by microglia (Iba1+) in the cortex. Astrocytes (GFAP+), hematopoietic cells (CD45+) and neuroblasts (Dcx+) do not express PK2 after TBI. Scale bar: 50μm.

Interestingly, we found that *Prok2* expression was upregulated in the injured cortex, beginning one day after the injury, with the highest levels achieved on the second day, and returning to basal level on day 7 (Fig. 4B). The analysis of the cortex tissue at the injury site showed that PROK2 was also upregulated at the protein level, and the PROK2-rich area was concentrated in the region surrounding the site of needle entry, inside the cells and in the extracellular space (Fig. 4C).

We next investigated which cells express PROK2 after brain injury. It has been previously reported that in cortical cultures PROK2 is mostly expressed in neurons, with a discrete expression in astrocytes and no expression in microglia. Under glutamate- and amyloid-beta-induced toxicity PROK2 is upregulated in both neurons and astrocytes (Cheng et al., 2012; Severini et al., 2015). Here we found that 7 days after the injury PROK2 was not expressed by astrocytes (GFAP+ cells) or immature neurons (Dcx+ cells) *in vivo* (Fig. 4D). Brain injuries cause a disruption of the blood-brain barrier turning the blood vessels permeable to blood-derived factors and cells (Gyoneva & Ransohoff, 2015). PROK2 is expressed by neutrophils and monocytes (Lecouter et al., 2004), and increases in PROK2 in neuropathologies have been associated to infiltrated cells (Abou-Hamdan et al., 2015; Giannini et al., 2009). With that in mind we investigated if hematopoietic cells that had infiltrated the brain parenchyma were the PROK2+ cells. We conducted an immunohistochemistry with PROK2 and CD45 antibodies. CD45, also known as leukocyte common antigen, is expressed in all nucleated hematopoietic cells, and was then used as a marker of these cells. No colocalization was found with the two antibodies (Fig. 4D). We next conducted an immunochistochemistry with PROK2 and Iba1 (a microglia marker) antibodies. Most of the PROK2+ cells were also Iba1+, demonstrating that microglia is the main source of PROK2. Despite the previous report of no expression of PROK2 by cultured microglia (Cheng et al., 2012), we show that its expression is only induced after brain injury, and it is not detected in the intact brain.

### PROK2 modulates the migration of SVZ cells to the injured cortex

Since PROK2 is important for the olfactory bulb neurogenesis acting as a chemotactic factor for SVZ cells, it is overexpressed in the injured cortex, and SVZ cells migrate to the cortex after the injury, we hypothesized that PROK2 could have an undescribed role in a pathological context, and act as a chemoattractant for SVZ cells to the injured area. To test this hypothesis we used a commercially available PROK2 antagonist, PKRA7, in mice submitted to brain injury. We administered BrdU for 2 days prior to the injury to label the SVZ cells, and PKRA7 (or DMSO in the control animals) was administered i.p. 1 hour after the injury and then every 24 hours for 3 days (Fig. 5A). The brains were analyzed on the fourth day. Fewer BrdU+ cells were found in the cortex of the PKRA7 treated mice compared to the DMSO treated group (Fig. 5B,C), suggesting that PROK2 signaling could be involved in the migration of SVZ cells to the injured cortex.

**Figure 5.**
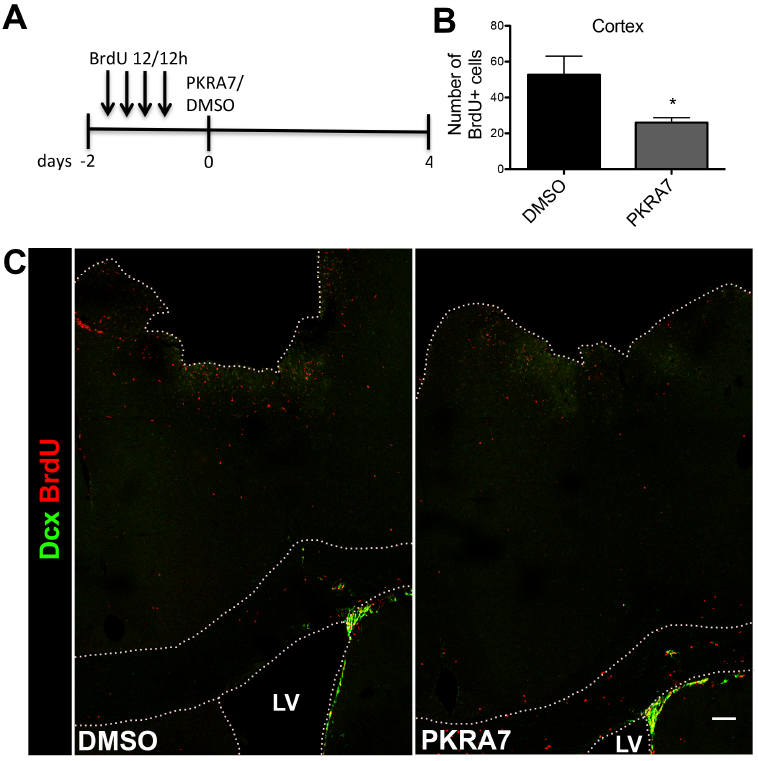
Inhibition of PK2 signaling decreases migration of SVZ-born cells to the injured cortex. (A) Scheme of BrdU and the PK2 antagonist PKRA7/DMSO administration. (B, C) Fewer BrdU+ cells (red) migrate to the cortex 4 days after TBI following administration of PKRA7. LV, lateral ventricle. Graphs show mean ± SEM. Scale bar: 100μm.

To address that, we used PROK2 expressing cells and a recombinant PROK2 in migration assays *in vitro* and *in vivo*. Co-culture experiments of spheroids of stable 293T cell line expressing mouse PROK2 and SVZ derived neurospheres were carried out in a collagen matrix for 5 days. When the neurospheres were co-cultured with control 293T cells the emigration of cells from the neurospheres were mainly symmetric, but when the neurospheres were cultured with the PROK2 expressing cells, the cells migrated longer distances from the neurospheres in direction to the spheroid (Fig. 6A). Similar results were found using cells expressing a human form of PROK2 and SVZ explants (Ng et al., 2005). We then injected recombinant PROK2 in the cortices of mice that were BrdU administered for 2 days (Fig. 6B). The same volume of normal saline was injected in control animals. On the fourth day the destination of the SVZ derived BrdU+ cells was analyzed. More than twice as many BrdU+ cells were found in the cortices of the PROK2 injected group mice (Fig. 6C, D, F). These data suggest that PROK2 is able to recruit SVZ cells to injured areas.

**Figure 6.**
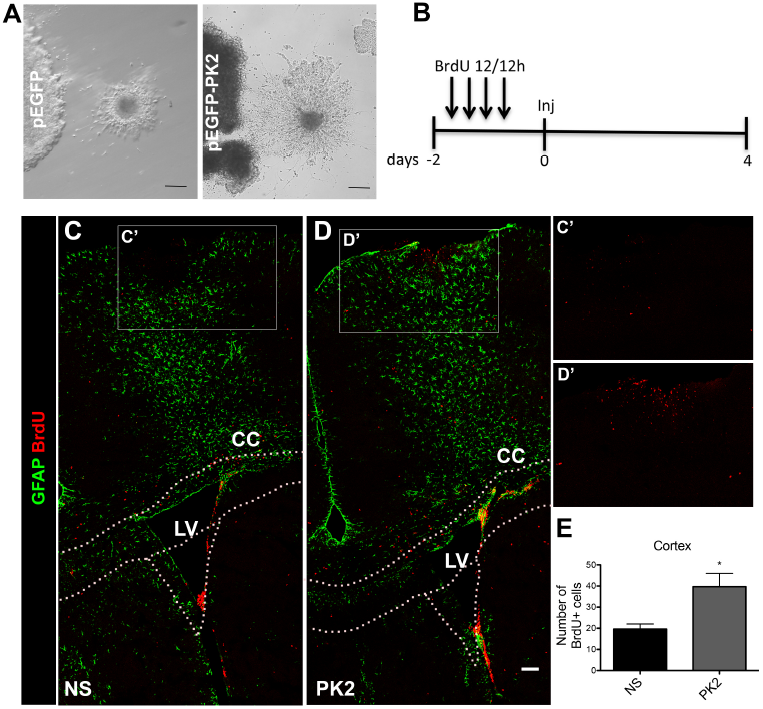
PK2 attracts SVZ cells *in vitro* and *in vivo*. (A) SVZ cells migrate longer distances when emigrate from the proximal side of the neurosphere in relation to cells that emigrate from the distal side. (B) Scheme of BrdU administration and injection of PK2 or saline. (C-E) More BrdU+ cells migrate to the cortex after injection of recombinant PK2. (C’, D’) Higher magnification of the cortices showed in C and D. LV, lateral ventricle; CC, corpus callosum; NS, normal saline. Graphs show mean ± SEM. Scale bar: 100μm.

Besides the physiological role of PROK2 as modulator of olfactory bulb neurogenesis (Ng et al., 2005), generation of circadian rhythms (Cheng et al., 2002; Li, J. D. et al., 2006), angiogenesis (Lecouter et al., 2001; Lecouter et al., 2003), reproduction (Maldonado-Perez et al., 2007) and gastrointestinal motility (Li, M. et al., 2001), PROK2 has recently been implicated in brain injuries and neurodegenerative diseases. In a mouse model of Parkinson disease PROK2 expression was induced in dopaminergic neurons and substantia nigra, promoting neuroprotection by activating ERK and Akt signaling pathways (Gordon et al., 2016). PROK2 was also showed to be an endangering mediator for cerebral damage in an ischemic model (Cheng et al., 2012), to modulate neuropathic pain in a chronic constriction injury model (Lattanzi et al., 2015), and to mediate amyloid-beta neurotoxicity in an Alzheimer’s disease model (Severini et al., 2015). To our knowledge this is the first report of cortical PROK2 induction after a traumatic brain injury.

The infiltration of neutrophils, lymphocytes and monocytes in the brain parenchyma after the initial injury triggers a wave of proinflammatory cytokines leading to the recruitment of immune cells and microglia (Lozano et al., 2015). Microglial activation induces the production and release of more proinflammatory cytokines and thereby increase the production of chemokines (Chodobski et al., 2011). PROK2 also upregulates a variety of chemokines (Curtis et al., 2013), and since chemokines are able to attract SVZ cells to injured areas (Filippo et al., 2013), PROK2 could indirectly promote chemoattraction of these cells by upregulating other chemokines, such as CXCL12. To test this hypothesis, we treated an astrocyte/microglia culture with recombinant PROK2 for 2 days and analyzed the expression of *Cxcl12* mRNA by qPCR. CXCL12 is a chemokine known to recruit SVZ cells to injured areas (Filippo et al., 2013). *Cxcl12* mRNA was not increased in the treated cells, but decreased instead (Fig. 7). *Prok2* mRNA was also not altered, indicating there was not a positive feedback loop of activation. *In vitro*, CXCL12 is only detected in astrocytes when they are activated (Dra. Marcella Reis personal communication). Indeed, we were not able to detect CXCL12 in the supernatant of astrocytes/microglia culture, even when they were treated with PROK2, measured by ELISA (data not shown). These data show that PROK2 was not able to induce the expression and secretion of CXCL12. We can not rule out that PROK2 induces the production of other chemokines that then recruit SVZ cells, but since we demonstrated that *in vitro* PROK2 can directly attract SVZ cells, it is possible that PROK2 act by a somatory of direct and indirect mechanisms.

**Figure 7.**
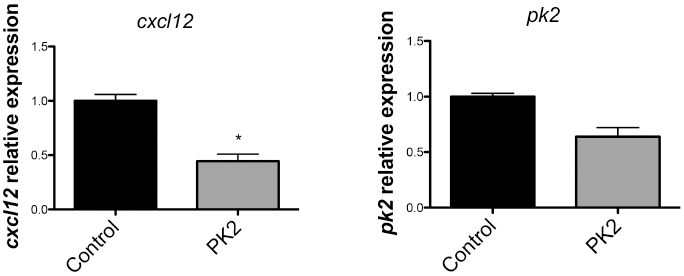
PK2 does not induce the expression of *pk2* or *cxcl12* in astrocytes and microglia *in vitro*. *cxcl12* and *pk2* expression by astrocytes and microglia in culture treated with recombinant PK2 for 2 days. Expression was normalized to *hprt*. Graphs show mean ± SEM.

Brain injuries trigger a cascade of pathophysiological processes that ultimately lead to cell death. The migration of SVZ cells to the injured area could help to replenish the cell population. PROK2 is a druggable molecule and could potentially be used as a modulator of SVZ cell migration, augmenting the amount of cells recruited to lesion areas, thus helping to promote a better recovery of acutely injured brain.

## Conclusion

We showed here that PROK2 is upregulated and expressed by microglia after cortical TBI in mice, and that PROK2 is able to recruit SVZ-derived cells to the site of injury. This is a new role described for PROK2 in the pathological context and it represents a potential target for stem cell-based therapies.

## Acknowledgements

Financial Support: This work was granted by FAPESP (Research Grants 2012/00652-5; 2012/06810-1) and CNPq (Research Grant 404646/2012-3; INCT-Regenera 465656/2014-5).

